# Task-generic and task-specific connectivity modulations in the ADHD brain: An integrated analysis across multiple tasks

**DOI:** 10.1101/755603

**Authors:** Roselyne J. Chauvin, Jan K. Buitelaar, Emma Sprooten, Marianne Oldehinkel, Barbara Franke, Catharina Hartman, Dirk J. Heslenfeld, Pieter J. Hoekstra, Jaap Oosterlaan, Christian F. Beckmann, Maarten Mennes

## Abstract

Attention-Deficit/Hyperactivity Disorder (ADHD) is associated with altered functioning in multiple cognitive domains and neural networks. This paper offers an overarching biological perspective across these. We applied a novel strategy that extracts functional connectivity modulations in the brain across one (P_single_), two (P_mix_) or three (P_all_) cognitive tasks and compared the pattern of modulations between participants with ADHD (n-89), unaffected siblings (n=93) and controls (n=84; total N=266; age range=8-27 years).

Participants with ADHD had significantly fewer P_all_ connections (modulated regardless of task), but significantly more task-specific (P_single_) connectivity modulations than the other groups. The amplitude of these P_single_ modulations was significantly higher in ADHD. Unaffected siblings showed a similar degree of P_all_ connectivity modulation as controls but a similar degree of P_single_ connectivity modulation as ADHD probands. P_all_ connections were strongly reproducible at the individual level in controls, but showed marked heterogeneity in both participants with ADHD and unaffected siblings.

The pattern of reduced task-generic and increased task-specific connectivity modulations in ADHD may be interpreted as reflecting a less efficient functional brain architecture due to a reduction in the ability to generalise processing pathways across multiple cognitive domains. The higher amplitude of unique task-specific connectivity modulations in ADHD may index a more “effortful” coping strategy. Unaffected siblings displayed a task connectivity profile in between that of controls and ADHD probands, supporting an endophenotype view. Our approach provides a new perspective on the core neural underpinnings of ADHD.

## Introduction

Attention-Deficit/Hyperactivity Disorder (ADHD) is a mostly early onset neurodevelopmental disorder characterised by symptoms of inattention and/or hyperactivity-impulsivity that are associated with impairments in multiple functional domains^1^. Multiple cognitive theories have been proposed to explain the underlying core deficits of the disorder, including a dysfunction in state and arousal regulation^2^, response inhibition^3^, broader executive functioning^4^, motivation^38^, and/or delay aversion^4^. Functional imaging studies building on these cognitive explanations have investigated the neural underpinnings of ADHD, but have revealed a heterogeneous pattern of altered neuronal function spread across the brain^1,5,6,37^. This fragmented pattern of findings asks for new integrated approaches that provide an overarching perspective on the functional architecture of the ADHD brain across cognitive domains.

Here, we offer such a perspective by applying a novel approach that integrates findings of cognitive tasks across multiple cognitive domains to assess the role of task-dependent connectivity modulations^7^. We thereby capitalize on the idea that task-induced connectivity patterns build on the baseline functional connectivity architecture as indexed by resting-state MRI analyses^8,9,10^. Using the resting-state architecture as baseline allows assessment of the regional specifics and magnitude of task-induced connectivity modulations across task paradigms^7^. Specifically, we use *task potency* as a metric to index the *strength* of task-induced connectivity modulations in terms of their difference from a resting-state baseline. This approach facilitates the comparison between task paradigms and permits to disentangle modulations that are shared across multiple cognitive functions (i.e. P_all_), resembling a cognitive core^11,12,13^, from those that are task-specific (i.e. P_single_). For example, a comparison of working memory, response inhibition, and reward tasks could reveal that participants with ADHD show alterations in the same network of inhibition-related brain regions in each task. This would provide support for theories that claim a prominent role for poor response inhibition in ADHD^3^. Alternatively, theories suggesting inefficient management of resources in participants with ADHD would be supported by observing, for example, a pattern of modulations that is highly specific to each task, while an overall cognitive core in support of task-general processes remains under-modulated^14,15^. Both theories are not incompatible: an alteration in one function network may induce coping strategies involving other functional networks; In combination, this may manifest as inefficiency across multiple task-generic and task-specific systems. In light of these possibilities we hypothesized that, in ADHD, the brain’s functional core interacts differently with more specialized network modulations, which could reflect inefficient use of the brain’s resources in ADHD.

To test our hypothesis, we applied our task-potency framework to a large cohort of participants with ADHD, their unaffected siblings, and healthy controls (N=266) and describe functional connectivity patterns across three cognitive domains (probing response inhibition^14^, working memory^390^, and reward processing^15^). Since unaffected siblings share on average 50% of genetic variation with their ADHD probands, the addition of unaffected siblings allowed assessing the impact of familial vulnerability - addressing the hypothesis that unaffected siblings show a task connectivity modulation profile that is an intermediate phenotype between that of diagnosed siblings and control participants^30, 43^.

## Methods

### Participants and (f)MRI acquisitions

We selected participants with ADHD, unaffected siblings of individuals with ADHD (but not related to the participants with ADHD included in this study), and typically developing controls (unrelated to any participant) from the NeuroIMAGE sample^16^. All selected participants completed an anatomical MRI scan, a resting state fMRI scan (RS), and at least one of the following task fMRI scans: a spatial working memory task (WM), a monetary-incentive-delay reward task (REWARD), and/or a stop signal response inhibition task (STOP) (see Table S2). Table 1 summarizes the demographics of the 89 participants with ADHD, 93 unaffected siblings, and 84 controls included in the current analyses. A full description of the selection criteria, task paradigms and MRI acquisition parameters is provided in Table S3 and supplemental method (appendices). Participants were scanned at two different sites; therefore we provide, in supplementary figures 7 and 8, a replication of all results within site and with matched samples on age, IQ, site, gender.

**Table 1:** Participant information: descriptive, clinical variables, and distribution of scan modalities, for each group sample and tasks: Resting state (RS), stop signal paradigm (STOP), reward processing (REWARD), Working Memory (WM). 1 combined symptoms from KSADS and Conners. 2 ratio of Amsterdam/Nijmegen scan localisation. For participant exclusion, see Table S1. **For more detail on IQ and gender representation and testing of differences, see Figure S1. Replication of analysis for differences in sample (scanner/site, gender, IQ) is available in Figures S7 and S8.

|  | N used in<br>final<br>analyses | Age<br>min - max | Age mean<br>(std) | % female** | Site<sup>2</sup> | Inattention<sup>1</sup><br>(std) | Hyperactivity<sup>1</sup><br>(std) | IQ**<br>(std) |
| --- | --- | --- | --- | --- | --- | --- | --- | --- |
| **Healthy control participants** |  |  |  |  |  |  |  |  |
| **RS** | 84 | 10.8 – 23.5 | 16.7 (3) | 50% | 65% | 0.4 (1.1) | 0.3 (0.8) | 107.2 (13.8) |
| **STOP** | 46 | 10.8 – 23.4 | 16.7 (3.1) | 58.7% | 50% | 0.3 (1.0) | 0.2 (0.6) | 106.1 (14.1) |
| **REWARD** | 46 | 12.7 – 23.5 | 16.6 (2.9) | 50% | 55% | 0.6 (1.5) | 0.3 (0.9) | 108.1 (12.8) |
| **WM** | 66 | 10.8 – 23.5 | 16.5 (3.1) | 50% | 80% | 0.3 (1.0) | 0.3 (0.8) | 105.8 (13.4) |
| **Unaffected Siblings** |  |  |  |  |  |  |  |  |
| **RS** | 93 | 7.7 – 28.1 | 16.9 (4) | 57% | 45% | 0.7 (1.4) | 0.6 (1.1) | 101.6 (14.9) |
| **STOP** | 55 | 7.7 – 27.3 | 17.1 (4.1) | 54.5% | 36% | 0.7 (1.7) | 0.7 (1.3) | 102.6 (16.3) |
| **REWARD** | 57 | 9.1 – 28.1 | 17.2 (3.7) | 57.9% | 49% | 0.6 (1.4) | 0.7 (1.2) | 100.1 (15.2) |
| **WM** | 44 | 7.7 – 28.1 | 16.2 (4.1) | 56.8% | 54% | 0.3 (1.0) | 0.3 (0.7) | 102.6 (13.7) |
| **ADHD participants** |  |  |  |  |  |  |  |  |
| **RS** | 89 | 8.5 – 24.5 | 17.4 (3.1) | 31.5% | 38% | 7.3 (1.9)* | 5.8 (2.3)* | 94.4 (14.5) |
| **STOP** | 49 | 11.1 – 24.5 | 17.7 (2.7) | 22.4% | 37% | 7.3 (1.5)* | 5.3 (2.3)* | 95.8 (14.7) |
| **REWARD** | 57 | 10.2 – 24.5 | 17.7 (3.1) | 35.1% | 32% | 7.0 (1.5)* | 6.1 (2.0)* | 97.0 (14.9) |
| **WM** | 51 | 10.2 – 24.2 | 17.2 (3.23) | 35.3% | 47% | 7.5 (1.3)* | 5.8 (2.3)* | 94.6 (14.3) |

### Task potency calculation

Our task potency approach is described in detail elsewhere ^7^. In brief, for each participant and each pre-processed RS, WM, REWARD, and STOP fMRI acquisition (see eMethod for pre-processing procedures) we defined functional connectivity matrices using 179 regions from a hierarchical whole-brain atlas^17^ (see Figure S2). We calculated connectivity as the normalized Fisher-*Z* partial correlation between the timeseries of each pair of regions in the atlas (see Supplement Method). To isolate connectivity changes induced by task modulation (WM, REWARD, STOP) from changes in the brain’s baseline architecture (RS), we standardized each individual-level pair-wise correlation obtained during task acquisition by subtracting the corresponding pair-wise correlation value calculated for the RS scan of that participant. This effectively allows comparing each connection in the task connectivity matrices in terms of its magnitude of deviation from that participant’s resting baseline^7^. We refer to this deviation as ‘task potency’.

For each task, we created group-level task potency matrices by averaging the individual-level potency matrices across all participants in each diagnostic group. Within these group-level matrices we selected those connections that were *sensitive* to task modulation, by thresholding each group-level task potency matrix. Negative and positive thresholds were defined using a two-tailed version ^7^ of mixture modelling thresholding ^18^. We used the most conservative limit across groups (as controls, ADHD, and Sibling groups were not equal size) in order to compare the same level of information and selected potency values exceeding this limit as being sensitive to task modulation.

To integrate results across task-paradigms, we subdivided these sensitive connections depending on their modulation by one or more of the tasks. In particular, we refer to connections that were modulated by one task only as P_single_, to connections that were modulated by more than one but not all tasks as P_mix_, and to connections that were modulated regardless of task as P_all_. We verified that the relative percentage of participants that performed multiple acquisitions was equivalent between groups to avoid a possible bias related to participant by task interactions in edge selection (see details in supplementary table 2).

### Group differences in task connection type

To assess whether ADHD was associated with a deviant distribution of task-induced modulations across the brain and across tasks, we compared the distribution of task-sensitive connections and relative P_single,_ P_mix_ and P_all_ connections across the three diagnostic groups. We compared the amount of sensitive connections between groups by indexing the percentage of connections included for each group relative to the total number of sensitive connections. We assessed between-group differences in the specificity of connections by obtaining for each group the percentage of connections per type relative to the total number of sensitive connections for that group. Finally, we assessed the ratio of connections uniquely modulated by each diagnostic group (unique connections) versus those connections that were also modulated by one or both of the other groups (shared connections).

To account for the heterogeneity in the population, we used a bootstrapping procedure to statistically infer group differences against an appropriate null distribution. Defining empirical null distributions specifically for each diagnostic group allows controlling for sensitivity-differences due to relative group sizes across tasks and across diagnosis. We sub-selected 80% of the study population, computed the different metrics of interest, i.e. true values (percentage of connections per label and amplitude of modulation), randomly relabelled the participants keeping group size equal and computed the same values for a random expectation. We perform this sub-selection and procedure 10000 times in order to build a null distribution. We tested for significant differences between the average observed values across 10000 sub-selections and the obtained null distribution (for further details see Supplementary Methods).

Using this procedure, we tested differences in the percentage of connections modulated across tasks (task-sensitive, P_all_, P_mix,_ P_single_ connections), and in the percentage of shared versus unique connections for each of the connection labels. To further assess whether differences in percentage of selected connections were associated with different amounts of modulation, we compare their average amplitude of modulation and tested values against the corresponding diagnostic group specific null distribution. P-values were assessed for significance using FDR correction across tests per group at q<0.05. Replication of values in light of possible confounder effects (medication, gender, scanner of acquisition, and comorbidity) are presented in Figure S4.

Finally, we assessed the stability versus the heterogeneity of task connection types in the different groups. To this end we used the 10000 extracted values from the 80% sample bootstrapping procedure and computed each connection’s selection rate at the group level across bootstraps and its associated shared selection rate between two groups. We computed these rates for sensitive, P_all_, P_mix_ and P_single_ connections. These group-level selection rates index how specific a selected connection is to one particular group by computing the difference in selection rate between groups for each connection. We can then display the *uniqueness (for a specific group)* versus the *sharedness* (across groups) of each connection. By comparing both rates, we can estimate which connections are uniquely and reproducibly selected in one group only, potentially representing idiosyncratic strategies to solve a task.

## Results

### Establishing an ADHD connectivity profile

Starting from the set of connections that yielded significant connectivity modulations across all participants, we compared the diagnostic groups in terms of the sensitivity and specificity of their connectome to task modulations. Figure 1 shows that participants with ADHD modulated a significantly smaller part of sensitive connections (mean 40.1% of all sensitive edges, sd=4.7, p<0.005) compared to the group-specific null distribution (50.6%, sd= 4.1), while the percentage of edges modulated by controls and siblings was not significantly different from random expectation (47.4%, sd=4.4 and 44.5%, sd=4.5, respectively).

**Figure 1:**
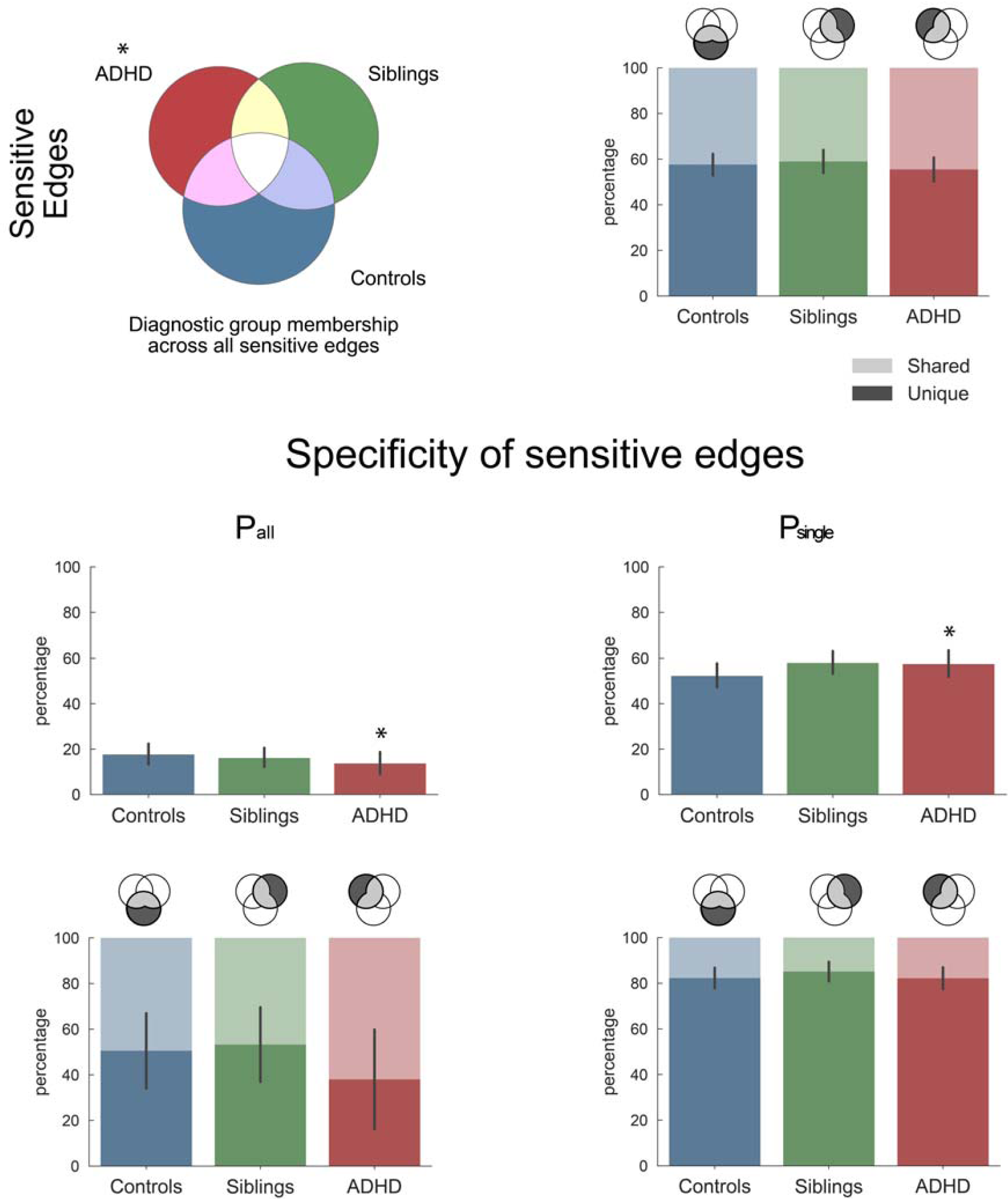
Sensitivity to task modulation. Description of total connectivity modulations across the three tasks and diagnostic groups (ADHD, siblings, controls). The first row shows for each group the percentage of connections they modulated across the three tasks (sensitive connections) and within these selected connections, the relative percentage of connections unique to one group or shared across groups. We further split the selected connections of each group into task-specific (P_single_), and common (P_all_) connections, corresponding to connections modulated in only one, or all three tasks, respectively. The second row of this figure quantifies the relative percentage of each connection type (P_all_ and P_single_) within the sensitive connections of each group. For the connections described in the second row, the third row then quantifies whether these connections were unique to that group or shared across groups. Stars indicate significant differences from null distribution after FDR correction (see table S5). Replication of these findings across possible confounding effects (scanner, gender, medication, comorbidity, age, IQ) is available in Figures S7 and S8.

Illustrating the task-specific nature of the sensitive connections, the bottom part of Figure 1 displays the proportion of selected connections that were P_all_ or P_single_ (full results including P_mix_ edges are available in Figure S4). ADHD had 13.8% P_all_ modulations, which was significantly lower than expected (p=0.031), and lower compared to both controls (17.8%) and siblings (16.3%). Within the P_all_ connections, ADHD seemed to mostly modulate connections that controls and siblings also modulated (the percentage of shared connections was more than 10% higher compared to the other groups, yet, this group difference did not reach significance).

In contrast to the lower number of P_all_ connections, the ADHD group exhibited a significantly higher percentage of P_single_ connections compared to the null distribution (see Figure 1). While the sibling group also showed a high percentage of P_single_ connections, this difference did not survive multiple comparison corrections across tests within this group. We observed no significant between-group differences in the uniqueness of the P_single_ connections.

### Reproducibility – from group level to individual analyses

We assessed the variability of the task connection types across bootstraps within each group, to examine the homogeneity of results across participants within a group. Figure 2 shows the results of these analyses for P_all_ and P_single_ edges (task-sensitive and P_mix_ related results can be found in FigureS5). The P_all_ connections in particular displayed strong homogeneity across control participants, with a notable shift toward 100% selected for controls, demonstrating a high reproducibility in controls and missing characterisation of these edges as P_all_ in other groups. In other words, controls reliably modulated P_all_ connections that were not consistently modulated by both other groups. In contrast, siblings seemed to modulate an alternative set of P_all_ connections that were not used by controls (figure 2 top right). This further informs on differences and similarities observed in Figure 1. In contrast, the P_single_ connections observed in ADHD and siblings (see Figure 1) were heterogeneous across participants, as illustrated by an absence of a shift in the distributions shown in Figure 2 towards the ADHD and sibling groups (see also Figure S5). Finally, we refer to Figure S6 for a description of the location of P_all_ connections that were strongly selected in each of the diagnostic groups.

**Figure 2:**
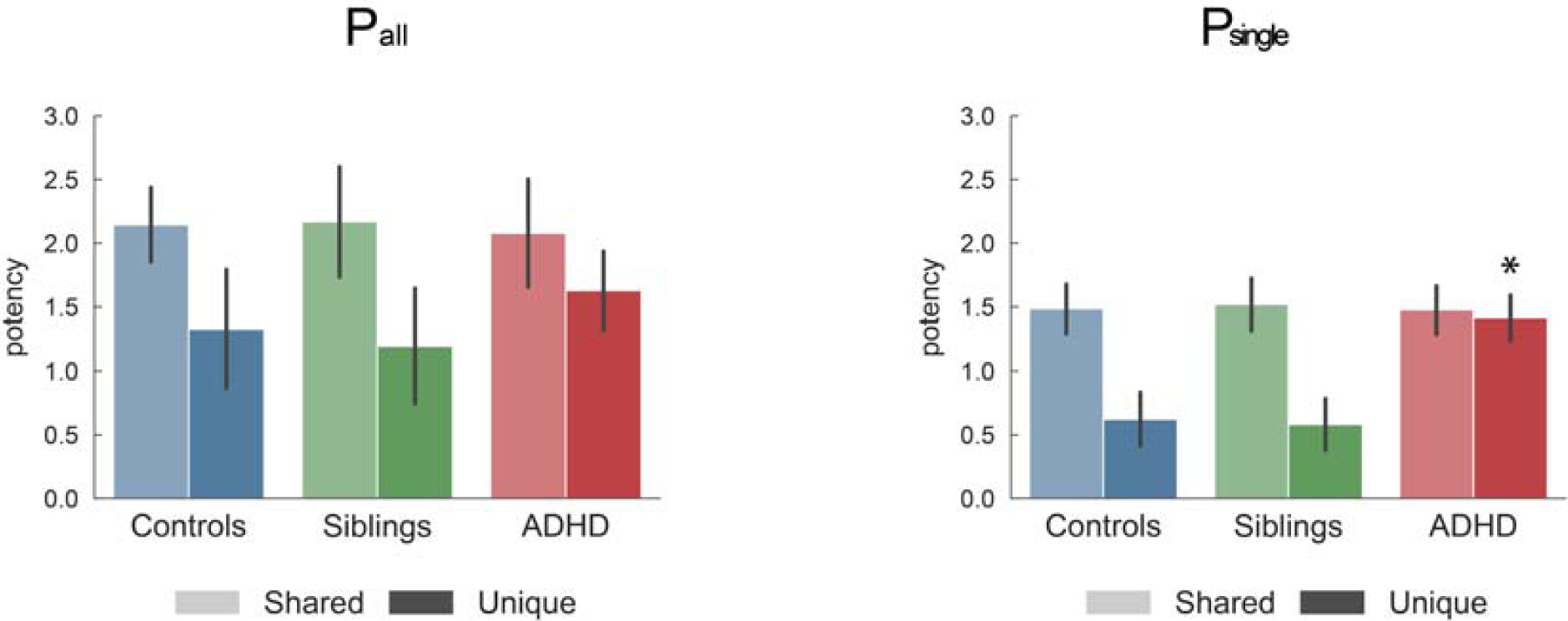
Comparison of selection reliability across bootstraps. By investigating the reproducibility of the selection of connections across bootstraps for each group and connections, we inferred on the uniqueness (x-axis) and shareability (y-axis) of each connection between two groups. Y-axis represents the percentage of bootstraps in which a connection is selected in two groups as a common or task specific connection, relatively. X-axis represents the difference between groups in percentage of selection across bootstraps of a connection as P_all_ or P_single_ edges. A connection that was always selected in both groups, i.e. high shareability, shown at the top corner of each triangle, would represent a connection that cannot be used to differentiate between those two groups. A connection that was always selected in one group only, located in the lower corners of the triangles, would be unique to a group and could be used to predict the group. Connections that would be heterogeneously selected in the population would have a low uniqueness (around 0 on the x-axis) and a low shareability (bottom of y-axis). The distribution at the basis of the triangle informs about the density of connections represented in the triangle, i.e. the spread of the distribution indicates whether only a small subset or a larger representation of connections are most often selected in one group relatively to the total amount of selected connections.

### Amplitude of modulation

As a proxy for cost estimation of different connectivity profiles reflecting possible compensatory mechanisms, we assessed whether the group differences in connectivity profiles were associated with group differences in the amplitude of the modulations. Figure 3 shows that siblings and controls equally modulated the different connection types (Table S4 provides statistical details). However, participants with ADHD overmodulated connections that were unique to them. This overmodulation was significant for P_single_ connections and P_mix_ connections (see Table S4), but did not reach significance for P_all_ connections. This is in line with the idea that the ADHD group placed more emphasis on P_single_ and P_mix_ rather than on generic connections.

**Figure 3:**
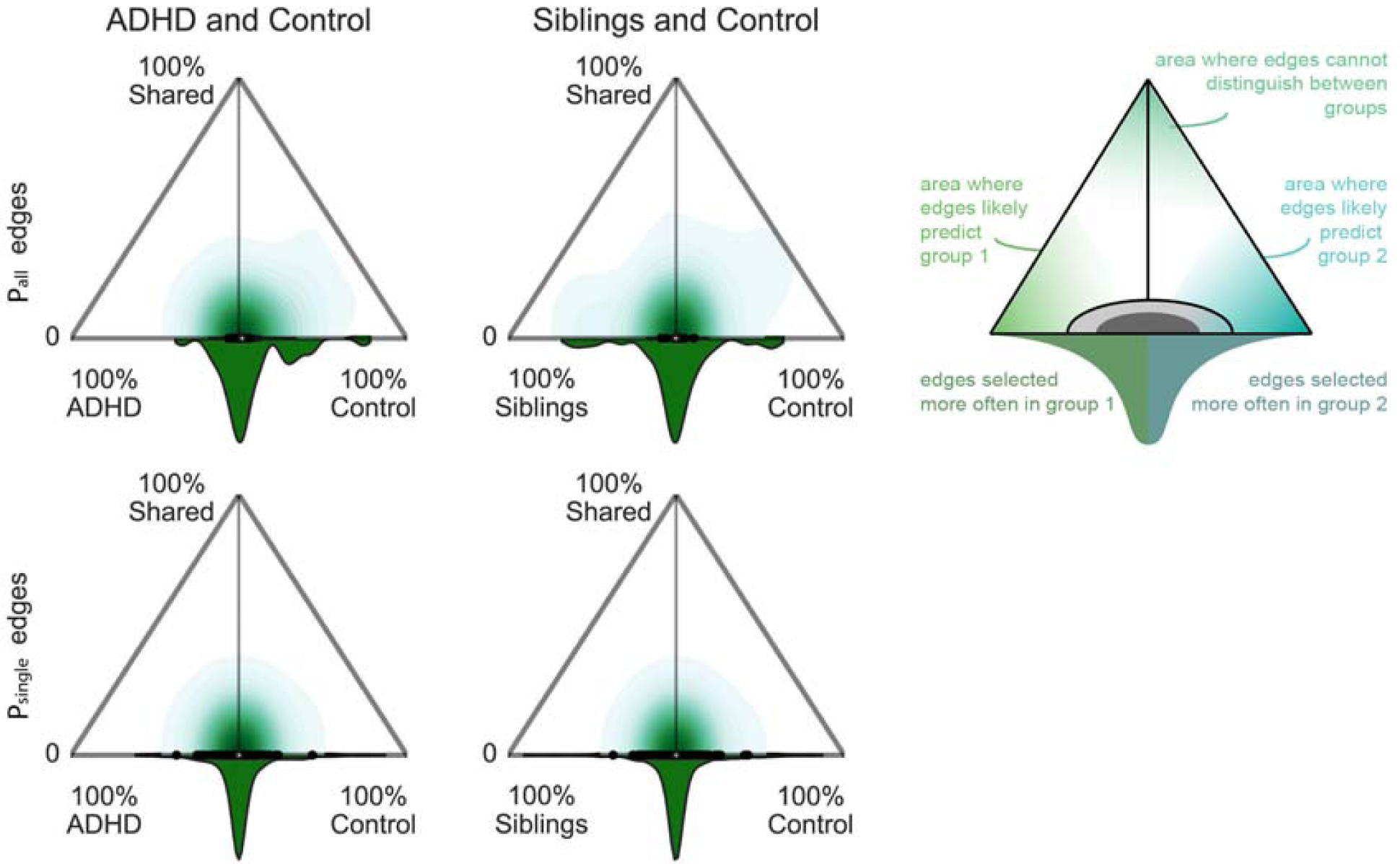
Modulation of edges. The graphs quantify the average task potency across unique or shared connections for each group and connection type (corresponding to the third line in figure 1). All reported values show the average and standard deviation across 10000 independent bootstraps. Indicated p-values show significant differences after FDR correction. Full results are available in Tbles S4-5. Replication of these findings in light of possible confounding effects (scanner, gender, medication, comorbidity, age, IQ) is available in Figures S7 and S8.

### Confounder Effects

In supplemental figures 7 and 8, we replicated figures 1 and 3 to investigate age, IQ, gender, comorbidity, scan site and medication effect with matched samples. The new sample size being reduced, we did not perform the bootstrapping procedure. We observed an overall replication of effects with a lower P_all_ and higher P_single_ percentage in ADHD and a partial similarity of connectivity profiles between unaffected siblings and ADHD groups.

## Discussion

We used a novel framework to provide an overarching perspective on the neurobiology of ADHD by inferring the nature of connectivity modulations in ADHD under the demands of working memory, reward processing, and response inhibition tasks. Our framework reveals that participants with ADHD activate significantly fewer connections than expected to perform each task. Furthermore, they modulated significantly fewer task-generic connections that share resources across tasks, and instead, relied significantly more on unique sets of connections (P_single_). In turn, participants with ADHD over-modulated those P_single_ connections, suggesting a task-tailored potential compensatory mechanism. In comparison, unaffected siblings of ADHD participants displayed an intermediate phenotype with values in between those observed for controls and ADHD.

The ADHD population is known for its clinical, biological and etiologic heterogeneity, and it is possible that different etiologic and/or biological mechanisms could result in the same behavioural symptoms. The results of our group-based analyses together with the high reproducibility at the individual level strongly support the idea that a core alteration underlies the cognitive and neural impairments observed in ADHD. Participants with ADHD rely less on a P_all_ core of modulations and instead involve task-tailored patterns of connectivity. These observations further support the idea that ADHD is characterized by neural inflexibility ^20,36^, as the use of predominantly task-tailored connectivity patterns makes task switching more demanding, more “expensive” and inefficient, with more challenging task performance as a result. As such, this connectivity profile provides support for the cognitive-energetic model of ADHD^2^. In this model, the limitations in arousal observed in ADHD could be a consequence of a higher level of energy required to perform cognitive tasks. This is potentially related to having to micro-manage P_single_ patterns instead of keeping a general processing core ready to perform. Assessing individual-specificities of these task-tailored connectivity in protocols including multiple tasks and mind-wandering detection could help disentangle whether this connectivity profile reflects coping strategies from possible distracting thoughts during the task.

Fitting with the hypothesis of inefficient neural processing is the observation that participants with ADHD typically are able to perform most tasks but exhibit large variability in the way they perform tasks^14,21,22^. Some studies have reported an inefficient use of resources for specific networks or functions in ADHD including the attention network^22,23^, executive functioning^245^, or cognitive control^22,25^. However, as these studies focus on specific cognitive aspects, they do not allow identifying a potential task-general underlying deficit, as demonstrated in the present study. Alternative approaches to investigating the efficiency of the brain’s organization have used graph theory and shown that the functional architecture of the ADHD brain is associated with differences in the balance of local and global efficiency^6,26-28^. However, these graph theory metrics provide no information on localized effects affecting specific cognitive functions. In contrast, our integrated approach provides a bridge between cognitive tasks and the functional architecture of the brain to understand the interaction between neural systems.

Unaffected siblings of individuals with ADHD share on average 50% of their genetic make-up with the ADHD probands, and accordingly, are hypothesized to share part of the ADHD endophenotype, i.e. biological deficits underpinning the ADHD phenotype ^29-31^. To avoid genetic relatedness between our groups, we selected unaffected siblings and ADHD from independent families, which reduced our sample size but validate a biological endophenotype. Siblings displayed a task connectivity profile in between that of controls and ADHD participants. Siblings showed a (non-significantly) larger use of P_single_ connections and some differences in the choice of P_all_ connections compared to controls. In terms of localization of P_all_ connections (see Figure S6), they combined motor connectivity like controls and subcortical connectivity like ADHD, which enables to perform the task as well as controls and which might be a more efficient alternative strategy, as it requires less modulation of connectivity (see Table S6). Previous research suggests that ADHD participants could compensate by using higher order executive systems or by relying more on lower-order visual, spatial, and motoric processing^32-35^. Our results support the literature by suggesting that unaffected siblings are potentially still able to recruit more task-general efficient connections.

The results shown in Figure 2 also highlight that P_single_ connections can be used to investigate such compensatory mechanisms at the level of individual participants. P_single_ connections are highly variable across participants, which is also described in previous work on task potency^7^. For instance, using longitudinal designs and models of compensatory strategies^35^, we can focus on those connections that are subject-specific and highly reproducible at the individual level to investigate a progressive specialization of individual compensatory mechanisms. In contrast to the high variability of P_single_ connections, the absence of some P_all_ connections was highly reproducible across ADHD and siblings compared to controls (Figure 2). As such, an absence of P_all_ connections may have potential as a biomarker for ADHD. This would encourage further, out-of-sample investigations of our new task potency approach.

Brain areas involved in the P_all_ connections and the associated differences between ADHD and controls are shown in Figure S6. At the brain regional level, participants with ADHD mainly missed modulations that connect regions within the executive control and reward pathways, including cerebellum, striatal, cingulum and cortical areas during task performance^26^. These results are coherent with meta-analysis showing hypoconnectivity in fronto-parieto-cingulo-striatal circuit^5,20^. As shown in Figure S6, participants with ADHD preserved only few P_all_ connections, interestingly involving striatal regions known to be involved in reward processing. Note that these results do not contradict typical findings of aberrant brain activity in reward-related regions in participants with ADHD^38^ as we showed that participants with ADHD used these connections with greater inconsistency and decreased modulation compared to controls. Knowing that ADHD participants make less efficient use of common pathways to govern multiple cognitive functions, will inform next studies aimed at understanding task response variability in ADHD.

Another important follow up is to integrate knowledge on resting state differences ^6^ As the task modulation builds upon the baseline architecture, after identifying differences between groups during task processing, both levels of baseline and modulation need to be integrated. This will also allow an understanding of the dependency between baseline and task effect and whether task connectivity differences can be predicted on the basis of baseline alterations^40^. The choice of task in this study is aimed to target cognitive functions altered in ADHD and follow up studies need to extend this finding to other cognitive domains to frame the P_all_ edges toward specific localized circuit or show a generalization of this effect. Additionally, longitudinal studies need to assess age-related trajectories of these findings, as we know that task potency changes with age^41^. The current study does not address a possible developmental delay explanation^42^.

In conclusion, we examined connectivity modulations across three tasks and demonstrated that individuals with ADHD showed less P_all_ and more P_single_ connectivity modulations when compared to controls or unaffected siblings. Our work represents an important step towards new integrative theories explaining how multiple neural alterations interact and result into multiple cognitive impairments in ADHD. Future studies should explore whether the results hold under other tasks paradigms.

## Acknowledgments

This work was supported by a Marie Curie International Incoming Fellowship under the European Union’s Seventh Framework Programme (FP7/ 2007–2013), Grant 327340 (Brain Fingerprint) to MM; The Netherlands Organization for Scientific Research (NWO) Grant NWO-Vidi 864-12-003 to CFB; and Wellcome Trust UK Strategic Award [098369/Z/12/Z] to CFB. Additional support was received from the European Community’s Horizon 2020 Programme (H2020/2014 – 2020) under grant agreements 667302 (CoCA) and 728018 (Eat2beNICE). BF is supported by a personal Vici grant of the Netherlands Organization for Scientific Research (NWO; grant 016-130-669). The NeuroIMAGE study was supported by NIH Grant R01MH62873, NWO Large Investment Grant 1750102007010 (to JB), ZonMW grant 60-60600-97-193, NWO grants 056-13-015 and 433-09-242, and matching grants from Radboud University Medical Center, University Medical Center Groningen and Accare, and Vrije Universiteit Amsterdam. The research leading to these results also received support from the European Community’s Seventh Framework Programme (FP7/2007-2013) under grant agreement number 278948 (TACTICS) and 603016 (MATRICS) and the Innovation Medicine Initiative grants 115300 (EU-AIMS) and 777394 (AIMS-2-TRIALS).

