## Supplemental Material for "Task-generic and task-specific connectivity modulations in the ADHD brain: An integrated analysis across multiple tasks"

#### Supplemental Appendix: Participant selection criteria

All subjects participated in the NeuroIMAGE project, the Dutch follow-up of the International Multicenter ADHD Genetics (IMAGE) study. Details about ethics approval, recruitment, assessment, and the general testing procedures can be found in the general methods and design paper of the NeuroIMAGE project (Rooij et al., 2015). In short, ADHD diagnosis was based on semi-structured interviews (the Schedule for Affective Disorders and Schizophrenia for SchoolAge Children [K-SADS] (C. Kaufman et al., 1997)) as well as the Conners ADHD questionnaires (Conners et al., 1998a,b). Probands with ADHD had to have six or more hyperactive/impulsive and/or inattentive symptoms according to DSM-IV criteria ("Psychiatry Online | DSM Library," n.d.); unaffected siblings and unrelated controls had to have less than two symptoms overall, based on a structured psychiatric interview (K-SADS) and Conners questionnaires. Inclusion criteria for MRI participation consisted of the absence of claustrophobia and any metal in the body. Informed consent was acquired from all participants, with parents supplying consent for participants less than 16 years old. All participants were required to not take their ADHD medication on the day of the testing, if they were under medication.

fMRI scans exhibiting limited brain coverage or excessive head motion were excluded from further processing. Limited brain coverage was defined as having less than 95% overlap with the MNI152 standard brain after registration of the fMRI scan to the MNI152 template. In addition, we excluded from each task those participants who were among top 5% in terms of head motion as quantified by RMS-FD, the root mean square of the frame-wise displacement, computed using MCFLIRT (Jenkinson, Bannister, Brady, & Smith, 2002). From this selection, we selected one participant per family to avoid enhancing similarity between or within groups.

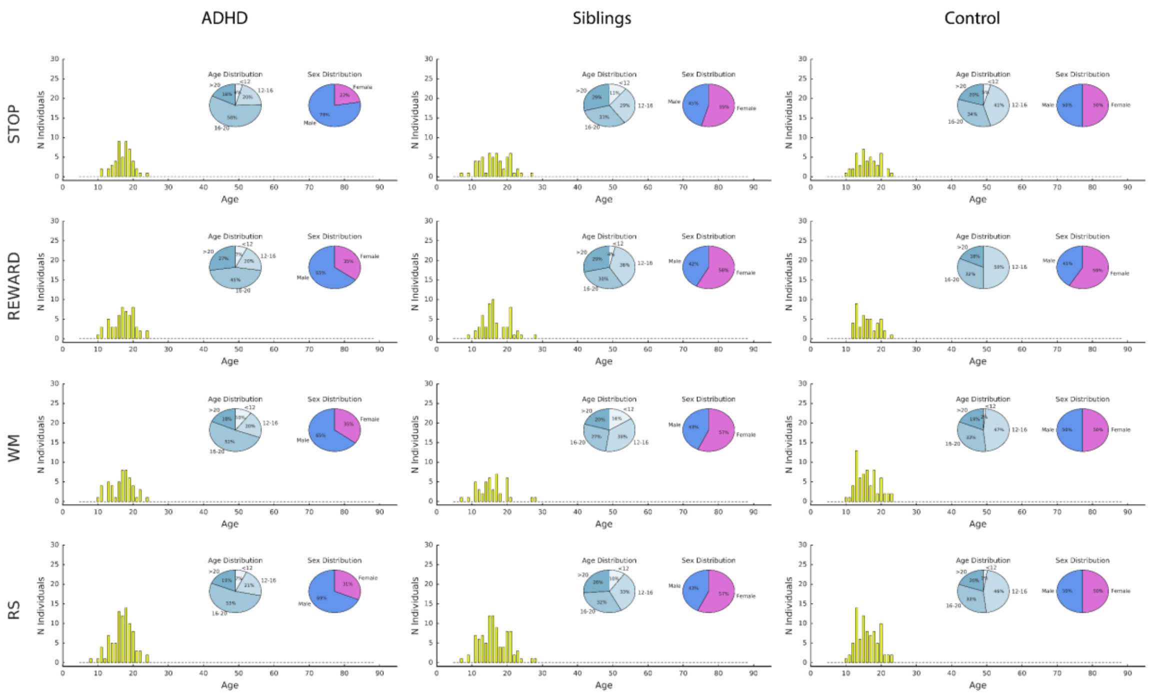


| Gender / IQ | ADHD | | | | Siblings | | | | Controls | | | |
|---|---|---|---|---|---|---|---|---|---|---|---|---|
| | WM | REWARD | STOP | RS | WM | REWARD | STOP | RS | WM | REWARD | STOP | RS |
| ADHD | | | | | 0.058 | **0.024** | **0.002** | **0.001** | 0.101 | **0.028** | **0.01** | **0.019** |
| Siblings | **0.008** | 0.284 | **0.032** | **0.001** | | | | | 0.612 | 0.905 | 0.798 | 0.43 |
| Controls | **$4x10^{-5}$** | **0.0002** | **0.001** | **$2x10^{-8}$** | 0.237 | **0.006** | 0.263 | **0.015** | | | | |

*Supplemental Figure 1: group and task population demographics. p-values of demographic differences between groups, upper triangle shows p-values from chi square test on gender, lower triangle shows p-values from independent t-test on IQ. P-values are uncorrected.*

*Supplemental Table 1: Overview of participant exclusion and final inclusion*

| | N initial with complete acquisition<br>without:<br>- Low T1 quality<br>- Incidental findings | N with both a quality RS and at least a task | insufficient MNI coverage | 5% highest mover rejection | N without family relationship **used in final analyses** |
|---|---|---|---|---|---|
| **Healthy control participants** | | | | | |
| **RS** | 136 | 130 | 2 | 6 | 84 |
| **STOP** | 131 | 67 | 2 | 6 | 46 |
| **REWARD** | 124 | 70 | 1 | 6 | 46 |
| **WM** | 153 | 99 | 2 | 7 | 66 |
| **Unaffected siblings of ADHD participants** | | | | | |
| **RS** | 112 | 107 | 1 | 5 | 93 |
| **STOP** | 106 | 64 | 2 | 5 | 55 |
| **REWARD** | 109 | 65 | 0 | 5 | 57 |
| **WM** | 85 | 51 | 1 | 4 | 44 |
| **ADHD participants** | | | | | |
| **RS** | 188 | 179 | 4 | 9 | 89 |
| **STOP** | 176 | 88 | 3 | 8 | 49 |
| **REWARD** | 197 | 116 | 3 | 9 | 57 |
| **WM** | 166 | 88 | 2 | 8 | 51 |

*Supplemental Table 2: number of participants having performed multiple tasks. in grey, in the diagonal is the total number of participants in final sample*

| | STOP | REWARD | WM |
|---|---|---|---|
| **Healthy control participants** | | | |
| STOP | 46 | 12 | 29 |
| REWARD | - | 46 | 32 |
| WM | - | - | 66 |
| **Unaffected siblings of ADHD participants** | | | |
| STOP | 55 | 23 | 19 |
| REWARD | - | 57 | 21 |
| WM | - | - | 44 |
| **ADHD participants** | | | |
| STOP | 49 | 22 | 17 |
| REWARD | - | 57 | 22 |
| WM | - | - | 51 |

No subject has performed all three tasks. Difference in representation of tasks overlap relative to the number of participants per group are not significant based on chi square tests (ADHD vs Siblings, p=0.99; ADHD vs Controls, p=0.99; Siblings vs Controls, p=0.99)

### Supplemental Appendix: Description of task-fMRI paradigms

#### *Stop signal task (STOP)*

A visual version of the stop signal task (Logan, Cowan, & Davis, 1984; van Rooij et al., 2015; von Rhein, Mennes, et al., 2015) was used to measure response inhibition during fMRI acquisition. In this task, participants had to respond as quickly as possible to a go-stimulus by left or right button press, unless shortly after presentation it was followed by a stop signal, in which case they were to withhold their response (25% of trials). The task consisted of two practice blocks and four test blocks, each consisting of 60 trials. For further details of the task and its acquisition parameters we refer to van Rooij et al., 2015.

#### *Monetary reward processing task (REWARD)*

A modified version of the MID task (Hoogman et al., 2011; Knutson, Fong, Adams, Varner, & Hommer, 2001; von Rhein, Cools, et al., 2015; von Rhein, Mennes, et al., 2015) was used to measure reward processing during fMRI acquisition. Participants were asked to respond as quickly as possible to a target by pressing a button. Prior to this target, a cue indicated the possibility to gain a reward after a button press within a given time window. Every trial ended with a feedback screen informing about the outcome of the current trial. Depending on the participants' performance, the response window for a correct response was adapted in the next trial resulting in an expected hit rate of 33%. The experiment lasted 12 minutes and a total of € 5 could be gained. For further details of the task and its acquisition parameters we refer to von Rhein et al., 2015.

#### *Spatial working memory (WM)*

The spatial span task used to measure spatial working memory is an adapted version of a task developed by Klingberg and colleagues (Klingberg, Forssberg, & Westerberg, 2002; McNab et al.,

2008; van Ewijk et al., 2015; von Rhein, Mennes, et al., 2015). Two trial types (baseline and working memory) and two memory loads (low and high) were implemented in the task. Each trial consisted of a sequence of either three or six yellow circles (low and high memory load, respectively), displayed on a 4×4 grid for 500 ms each, with a 500 ms inter-stimulus interval in between. Subsequently, during a 2000 ms response window, a probe consisting of a number with a question mark was presented in one of the 16 locations. During working memory trials, participants were asked to remember the spatial location and temporal order of the presentation of cues, and indicate with a 'yes' or 'no' response (left or right button, respectively) whether the location of the probe had been stimulated before, at the indicated temporal position. During baseline trials, red circles followed by the probe (always the number 8) were presented sequentially in the four corners of the grid in a predictive manner, and participants were required to pay attention but not to try to remember the sequence, and always had to press the 'no' button. During both conditions, feedback was presented after the response in the form of a green or red coloured bar below the probe (for correct and incorrect responses, respectively), for the remainder of the response window. The task was administered in four blocks of 24 trials each (presented in fixed random order), with a short break in between blocks to motivate participants and to avoid fatigue effects, with a total task duration of approximately 16 min. For further details of the task and its acquisition parameters we refer to van Ewijk et al., 2015.

*Supplemental Table 3: fMRI acquisition parameters and characteristics*

| | RS | STOP | REWARD | WM |
|---|---|---|---|---|
| **Image acquisition parameters** | | | | |
| General parameters | TE=40 ms, FOV=224mm, 37 axial slices, flip angle=80, matrix size=64x64, in-plane resolution=3.5mm, slice thickness/gap=3.0mm/0.5mm | | | |
| N volumes | >260 | 86 * 4 blocks | >300 | 107 * 4 blocks |
| N first volumes rejected* | 5 | 4 | 5 | 3 |
| **Healthy control participants** | | | | |
| N used in final analyses | 84 | 46 | 46 | 66 |
| TR (ms) | 1960** | 2340** | 2340** | 2340 |
| RMS-FD min-max | 0.031 - 1.218 | 0.029 - 0.360 | 0.032 - 0.408 | 0.033 - 1.328 |
| RMS-FD mean (std) | 0.138 (0.171) | 0.091 (0.066) | 0.130 (0.093) | 0.155 (0.193) |
| N ICs extracted – mean (std) | 35.32 (8.87) | 25.98 (3.54) | 65.09 (11.58) | 31.16 (5.32) |
| NICs defined as noise – mean (std) | 11.89 (4.06) | 7.79 (2.06) | 21.70 (5.24) | 9.77 (2.56) |
| **Unaffected siblings of ADHD participants** | | | | |
| N used in final analyses | 93 | 55 | 57 | 44 |
| TR (ms) | 1960** | 2340** | 2340** | 2340 |
| RMS-FD min-max | 0.026 - 1.357 | 0.030 - 0.947 | 0.039 - 0.331 | 0.029 - 1.504 |
| RMS-FD mean (std) | 0.159 (0.206) | 0.107 (0.140) | 0.124 (0.084) | 0.199 (0.293) |
| N ICs extracted – mean (std) | 36.56 (9.82) | 26.33 (4.41) | 61.61 (11.34) | 31.41 (5.18) |
| NICs defined as noise – mean (std) | 12.03 (4.41) | 7.87 (2.02) | 20.75 (5.10) | 8.94 (2.13) |
| **ADHD participants** | | | | |
| N used in final analyses | 89 | 49 | 57 | 51 |
| TR (ms) | 1960** | 2340** | 2340** | 2340 |
| RMS-FD min-max | 0.033 – 3.189 | 0.027 - 0.478 | 0.035 – 1.153 | 0.048 – 0.834 |
| RMS-FD mean (std) | 0.239 (0.381) | 0.129 (0.113) | 0.173 (0.181) | 0.259 (0.218) |
| N ICs extracted – mean (std) | 40.44 (9.37) | 26.78 (4.43) | 64.43 (13.72) | 34.13 (4.82) |
| NICs defined as noise – mean (std) | 12.31 (3.69) | 7.93 (1.82) | 21.38 (5.62) | 9.44 (2.20) |

* The number of initial volumes removed from further analyses varied to ensure comparability with earlier studies that used these data. Note that this variation will have very limited impact on the current analyses.

** some subjects were scanned with a different TR. In siblings: RS - 1860 for 2 subject; STOP – 2150 for 2; REWARD – 2280 for 1, in ADHD participants: RS - 1860 for 8 subjects; STOP – 2150 for 6 and 2280 for 1; REWARD – 2150 for 6 and 2280 for 1.

#### Supplemental Appendix: MRI preprocessing

All fMRI acquisitions were processed using tools from FSL 5.0.6. (FSL; http://www.fmrib.ox.ac.uk/fsl) (Mark Jenkinson, Beckmann, Behrens, Woolrich, & Smith, 2012; Smith et al., 2004; Woolrich et al., 2009). We employed the following pipeline: removal of the first 4 or 5 volumes to allow magnetization equilibration (see Supplemental Table 1), head movement correction by volume-realignment to the middle volume using MCFLIRT, global 4D mean intensity normalization, and 6mm FWHM smoothing. We denoised all preprocessed data for secondary motion-related artefacts using ICA-AROMA (Pruim, Mennes, Buitelaar, & Beckmann, 2015; Pruim, Mennes, van Rooij, et al., 2015) (Pruim 2015a, Pruim 2015b). Finally, we regressed out signal from CSF and white matter, and applied a 0.01Hz high-pass filter.

For each participant, all acquisitions were registered to its high-resolution T1 anatomical image using Boundary-Based Registration (BBR) available in FSL FLIRT (M. Jenkinson & Smith, 2001; Mark Jenkinson, Bannister, Brady, & Smith, 2002). All high-resolution T1 images were registered to MNI152 space using 12-dof linear registration available in FLIRT and further refined using non-linear registration available in FSL FNIRT (Andersson, Jenkinson, Smith, & others, 2007). We used the inverse of the obtained transformations to bring a brain atlas to each participant's native space, and performed all further analyses in participant native space.

#### Supplemental Appendix: Brain atlas used for computing connectivity

For each functional imaging scan we defined connectivity matrices using regions defined in a hierarchical whole-brain functional atlas (van Oort et al., 2017). This atlas contains 185 non-overlapping regions and was defined through Instantaneous Correlation Parcelation (ICP) applied to resting state fMRI data of 100 participants of the Human Connectome Project (HCP; (Glasser et al., 2013)). In short, ICP aims to parcel larger regions into subregions based on signal homogeneity, where the optimal number of subregions is determined based on split-half reproducibility at the cohort level.

Supplemental Figure 2 illustrates the hierarchical brain atlas, where areas were grouped into 11 higher-level networks: 9 resting state networks (visual1, visual2, motor, right attention, left attention, auditory, default mode network (DMN), fronto-temporal and striatum), and 2 anatomical structures (subcortical areas, and cerebellum). These higher-level networks respectively contained 19, 12, 22, 22, 18, 8, 18, 13, 7, 23, and 23 subregions.

All analyses were performed in each participant's native space. To this end we transformed the atlas to each participant's native space using the inverse of the anatomical to MNI152 non-linear warp, and the inverse of the linear transformation of the functional image to the participant's high resolution anatomical image. Voxel-membership in brain parcels was established on the basis of majority overlap. Areas that were on average across our population over 50% outside of the brain were rejected from further analyses. This resulted in the rejection of one area in brainstem. For consistency, we removed the 5 others brainstem areas. As a result, we used 179 areas to compute connectivity matrices, as explained below.

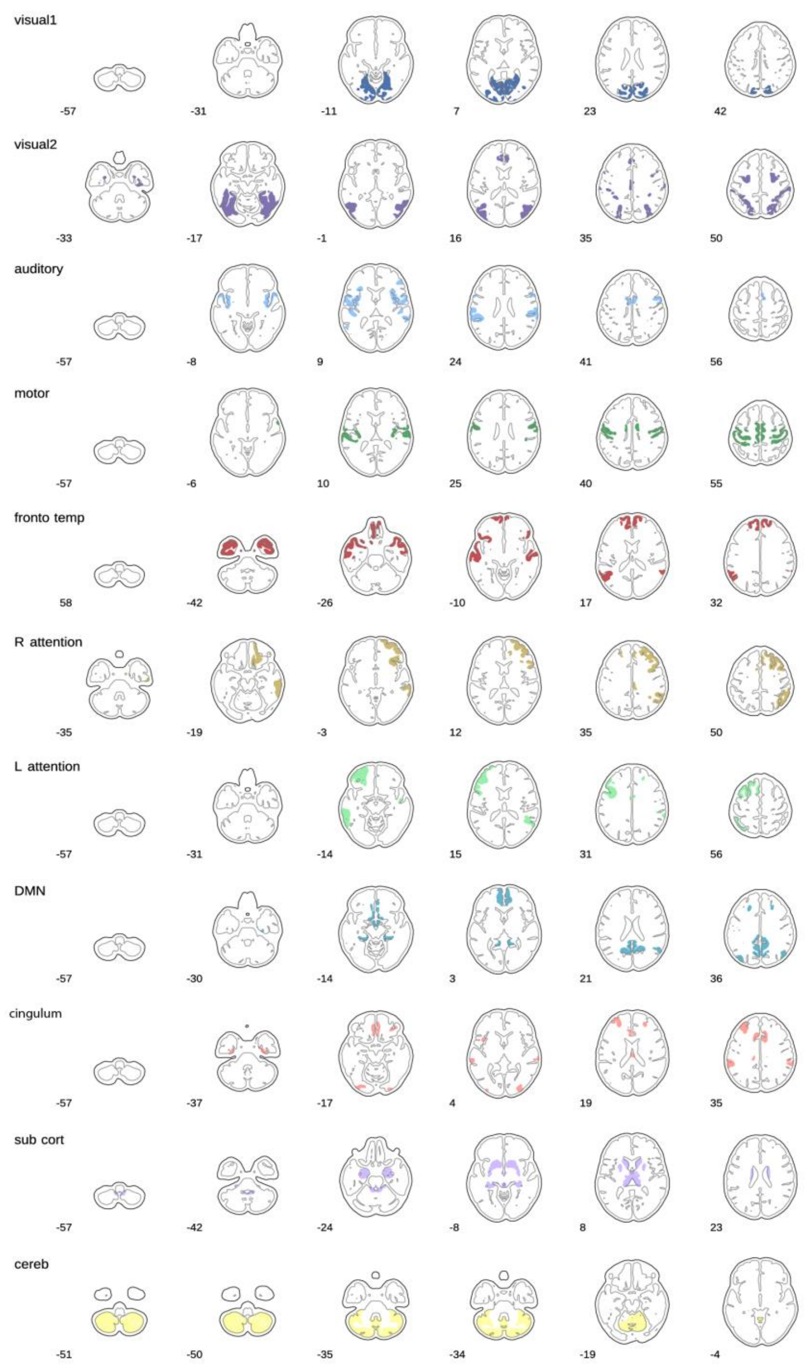


*Supplemental Figure 2: 179 areas selected from an ICP-based parcellation of the human brain (van Oort et al). Each area is coloured in accordance to its overarching network. Eleven large-scale networks constitute the first level of the parcellation: visual 1, visual 2, auditory, motor, fronto-temporal (fronto temp), right and left attention (R_attention, L_attention, respectively), default mode (DMN), cingulum, sub-cortical (sub cort), cerebellum (cereb) networks. We used the 179 regions that are part of the sub-network scale parcellation to obtain functional fingerprints based on 179x179 correlation matrices.*

#### Supplemental Appendix: Task Potency Calculation

To compute the partial correlation for each pair of regions we obtained each region's time series through multivariate spatial regression, using all 179 regions as regressors and each task's preprocessed full acquisition time series as dependent variable. The resulting regional time series were demeaned. For the WM and STOP task we temporally concatenated the time series of individual runs. Using these time series, we calculated 179x179 partial correlation matrices through inverting covariance matrices estimated by the Ledoit-Wolf normalization algorithm (Ledoit & Wolf, 2004) as implemented in nilearn (http://nilearn.github.io/). Finally, all pair-wise correlations were Fisher *r*-to-*Z* transformed.

To allow comparing connectivity values between acquisitions, we normalized the connectivity values within each matrix to fit a Gaussian distribution (Supplemental Figure 3). Importantly, we were cautious not to affect the tails of the connectivity distributions as these represent the most interesting connectivity values. Therefore, we modelled the obtained connectivity values per task using a Gaussian-gamma mixture-model to obtain "mixture-model-corrected" Z-stat values (Feinberg et al., 2010; Llera, Vidaurre, Pruim, & Beckmann, 2016). This model fits three curves to represent the data: a central Gaussian distribution representing the noise and two gamma distributions on each side of the central Gaussian that represent the signal as the tails of the data distribution. We used the main Gaussian, i.e., the one fitting the body of the distribution, to normalize our connectivity values with respect to its main distribution (i.e., noise), while not taking into account the extremes (i.e., signal). In practice, we applied the mixture modelling to the upper triangle values of each connectivity matrix and subsequently normalized the connectivity values by subtracting the mean and dividing by the standard deviation of the obtained central Gaussian model. As a result, the values within the normalized, *Z*-transformed partial correlation matrices are readily comparable across tasks (Feinberg et al. 2010).

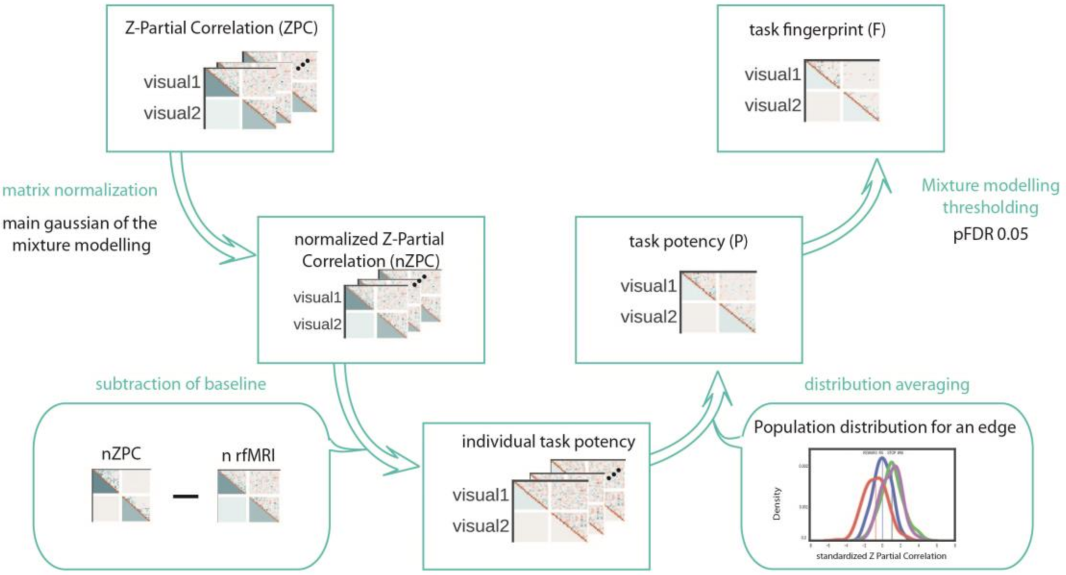


*Supplemental Figure 3: Task-potency pipeline. Using the brain parcellation shown in Supplemental Figure 2, we calculated 179x179 connectivity matrices for each individual in each task (WM, REWARD, STOP, RS). From the Fisher r-to-Z transformed partial correlation, we obtained task potency by first normalizing the task and rest connectivity and subsequently subtracting the rest from the task connectivity. Through population averaging and thresholding the resulting matrices we obtained a task potency fingerprint for each task (WM, REWARD, STOP).*

#### Supplemental Appendix: Permutation testing in selectivity

The reproducibility and significance of the group differences in task fingerprint is estimated by doing a permutation testing of the fingerprint similarity. To this aim, we repeated the analysis 10.000 times using 80% of each group, randomly selected and picked with replacement. Using the 10.000 bootstraps we obtained confidence intervals around each group's percentages in main Figure 1.

#### Subtype of edges analysis

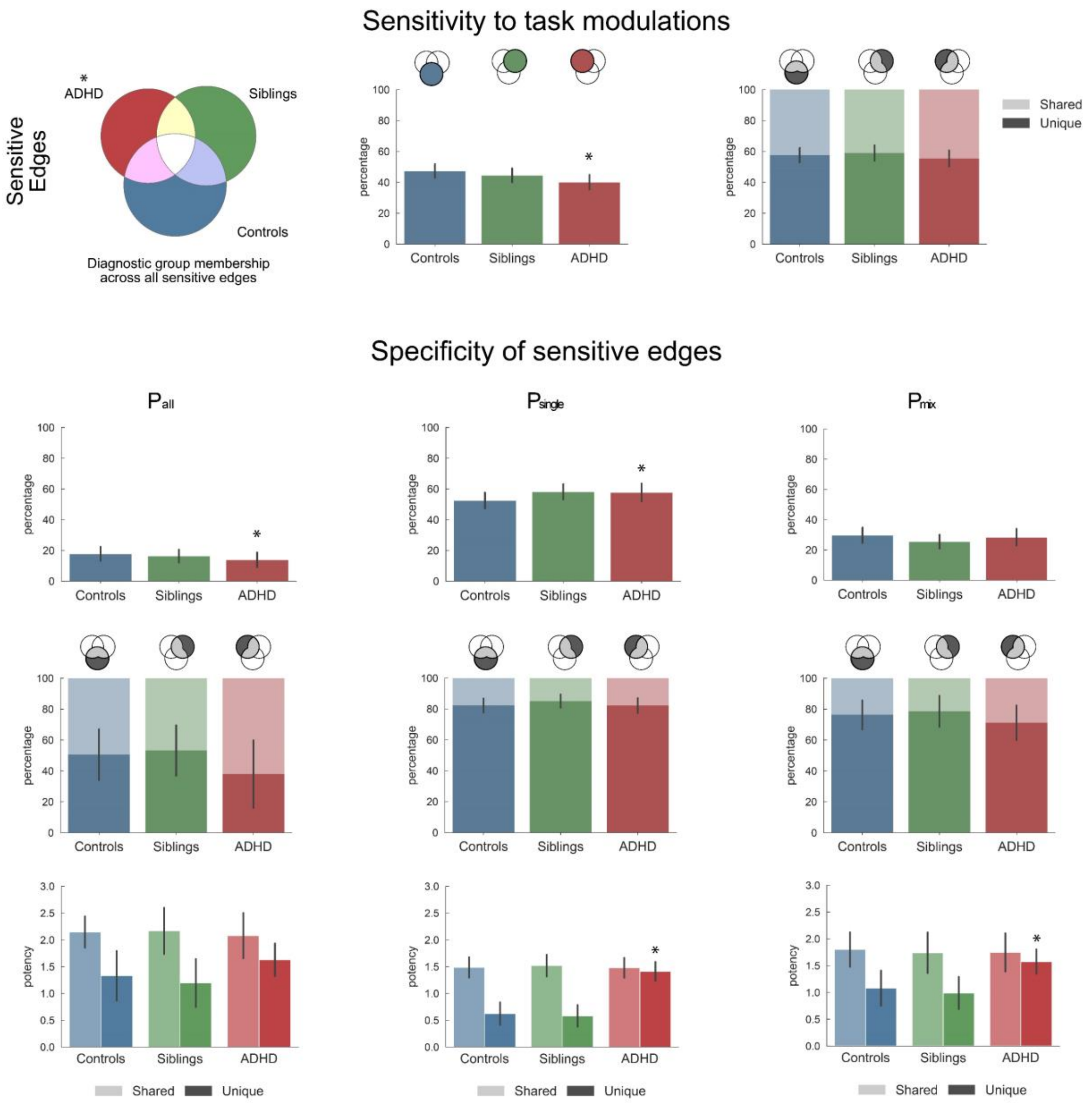


***Supplemental figure 4**: Description of connectivity modulations across the three tasks and diagnostic groups (ADHD, siblings, controls). This figure is an expansion of Figures 1 and 3 in the main text. The first row shows for each group the percentage of connections they modulated across the three tasks (sensitive connections) and within these selected connections, the percentage of connections unique to one group or shared across groups. We further split the selected connections of each group into $P_{single}$ , $P_{mix}$ and $P_{all}$ connections, corresponding to connections modulated in only one, two, or all three tasks, respectively. The second row quantifies the relative percentage of each connection type within the sensitive connections of each group. For the connections described in the second row, the third row then quantifies whether these connections were unique to that group or shared across groups. Finally, the fourth row quantifies the average task potency across unique or shared connections for each group and connection type. All reported values show the average and standard deviation across 10000 independent bootstraps. Indicated p-values shown significant differences after FDR correction for test within group. Full ANOVA results for the task potency results shown in row 4 are available in ST4-5. Replication of this findings for possible confounder effect (scanner, gender, medication, comorbidity) is available in SF7 and matching sample replication in SF8.*

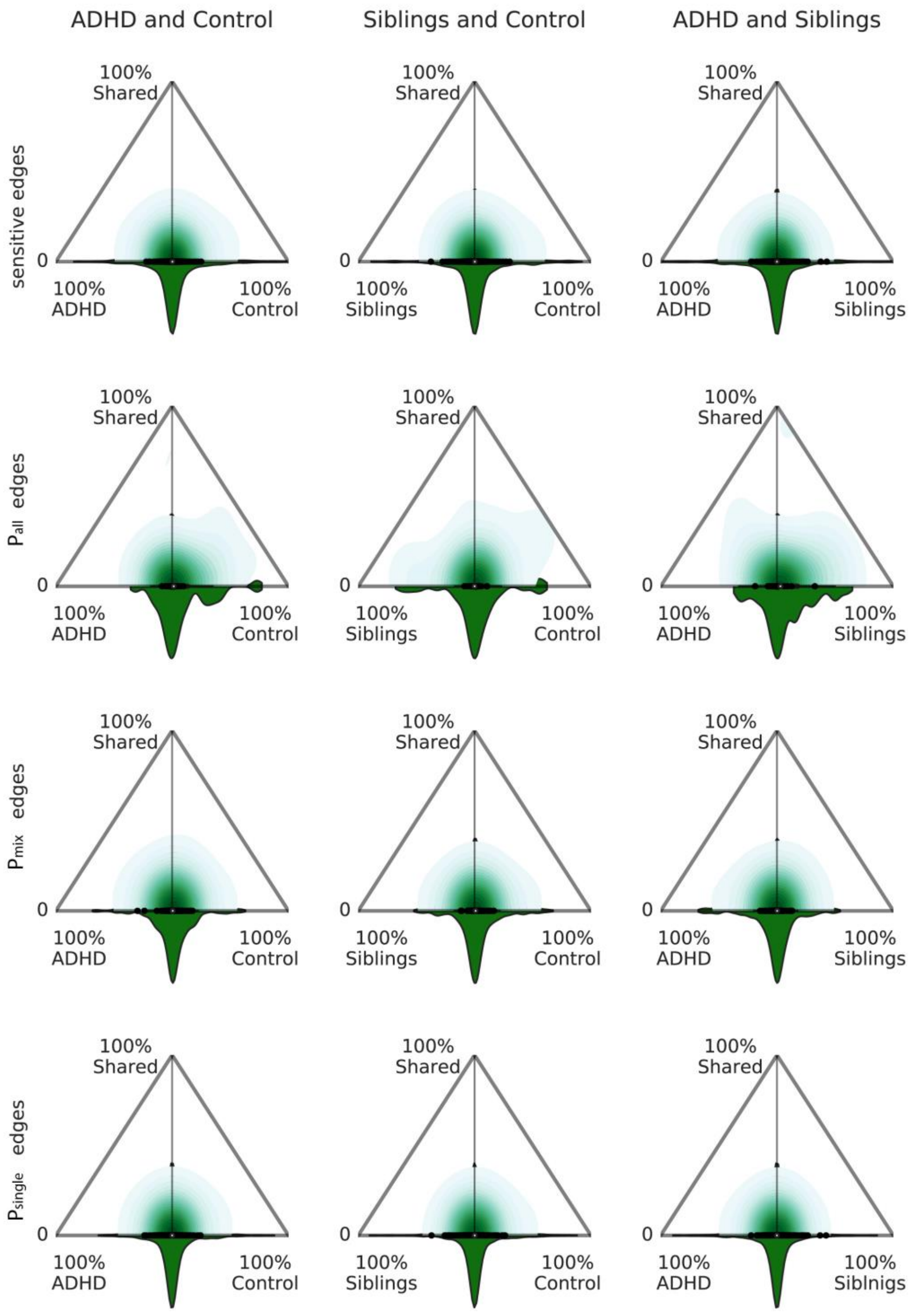


***Supplemental figure 5:*** *Comparison of selection reliability across bootstraps. This figure is an expansion of Figure 2 in the main text. By investigating the reproducibility of the selection of connections across bootstraps we inferred on the uniqueness (x-axis) and shareability (y-axis) of each connection between two groups. A connection that was always selected in both groups, shown at the top corner of each triangle, would represent a connection that cannot be used to differentiate between those two groups. A connection that was always selected in one group only, located in the lower corners of the triangles, would be unique to a group and could be used to predict the group. Connections that would be heterogeneously selected in the population would have a low uniqueness (around 0 on the x-axis) and a low shareability (bottom of y-axis). The distribution at the basis of the triangle informs about the density of connections represented in the triangle, i.e. the spread of the distribution indicates whether only a small subset or a larger representation of connections are most often selected in one group relatively to the total amount of selected connections. Random labelling corresponds to a random selection of participants while keeping percentages of participants across groups.*

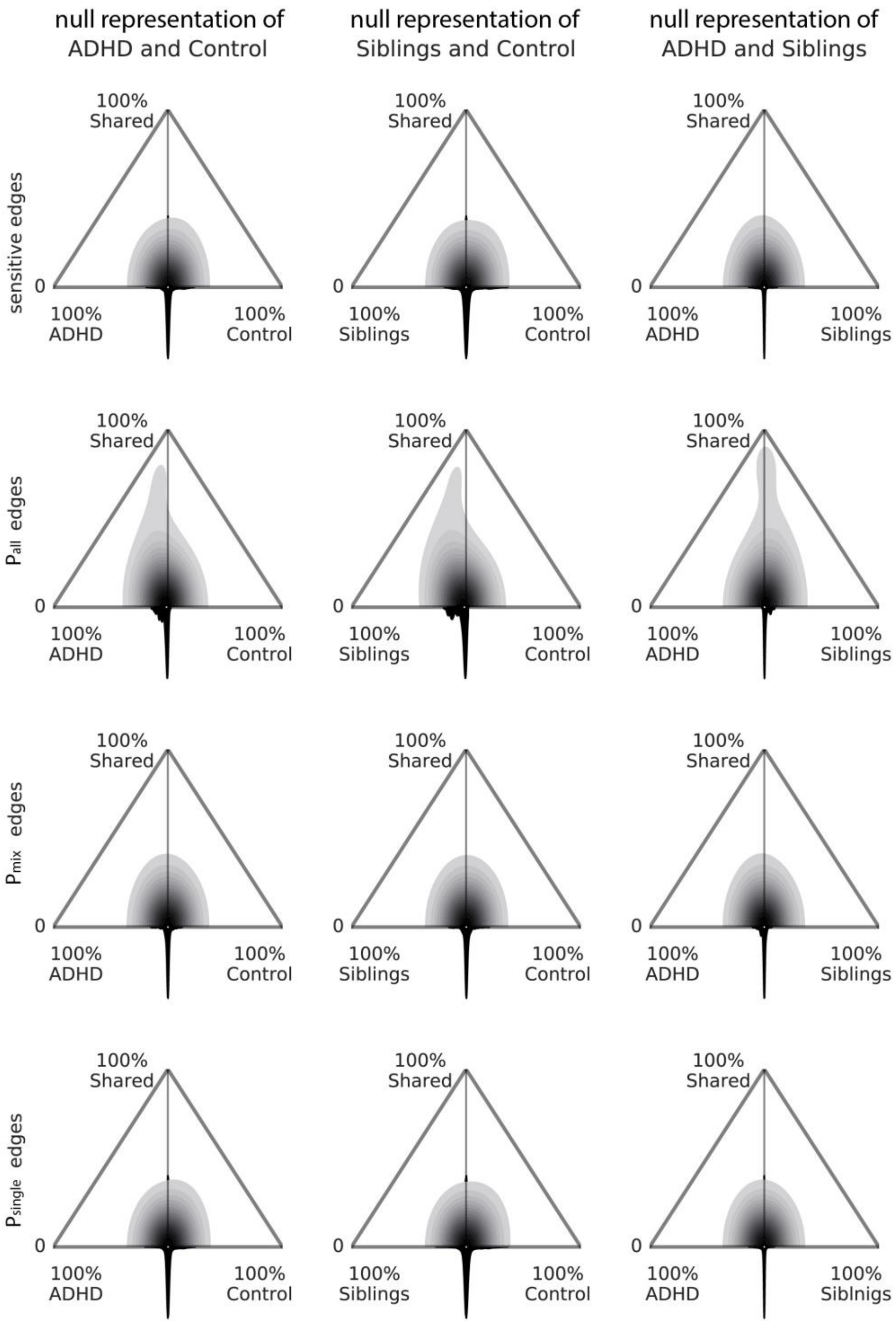


***Supplemental figure 5bis:*** *Comparison of selection reliability across bootstraps for the group corresponding null distribution.*

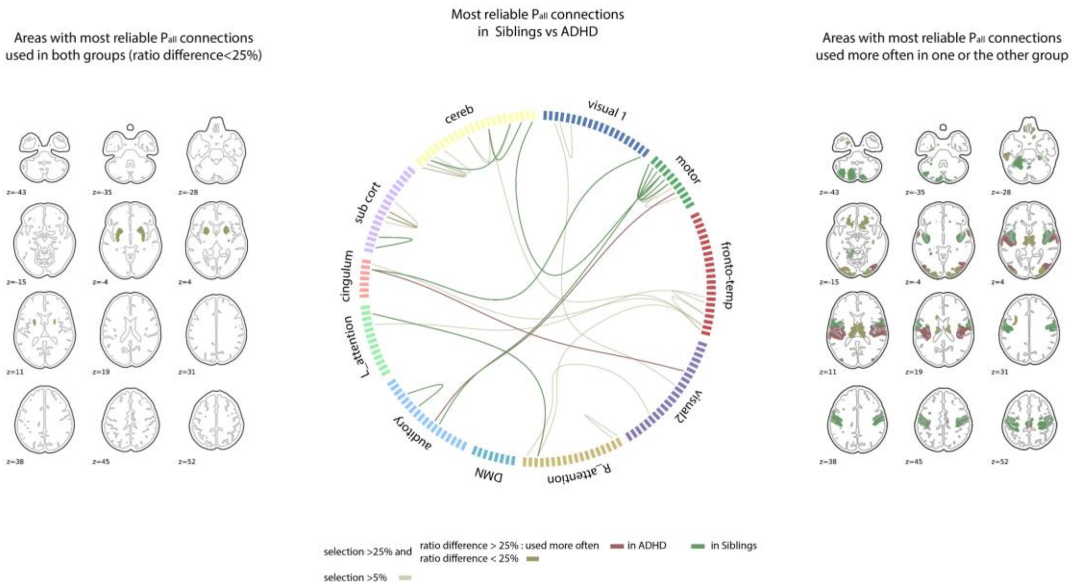
Areas with most reliable Pall connections used in both groups (ratio difference<25%)
Most reliable Pall connections in Siblings vs ADHD
Areas with most reliable Pall connections used more often in one or the other group
cereb
visual 1
motor
fronto-temp
visual2
R_attention
DMN
auditory
L_attention
cingulum
sub cort
selection >25% and
ratio difference > 25% : used more often
in ADHD
in Siblings
ratio difference < 25%
selection >5%

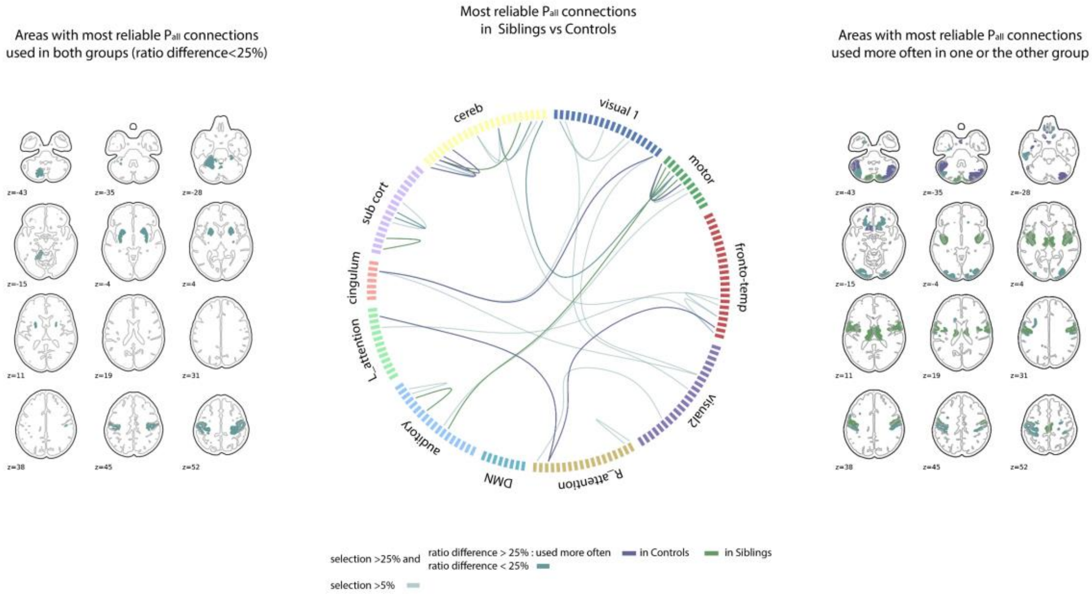
Areas with most reliable Pall connections used in both groups (ratio difference<25%)
Most reliable Pall connections in Siblings vs Controls
Areas with most reliable Pall connections used more often in one or the other group
cereb
visual 1
motor
fronto-temp
visual2
R_attention
DMN
auditory
L_attention
cingulum
sub cort
selection >25% and
ratio difference > 25% : used more often
in Controls
in Siblings
ratio difference < 25%
selection >5%

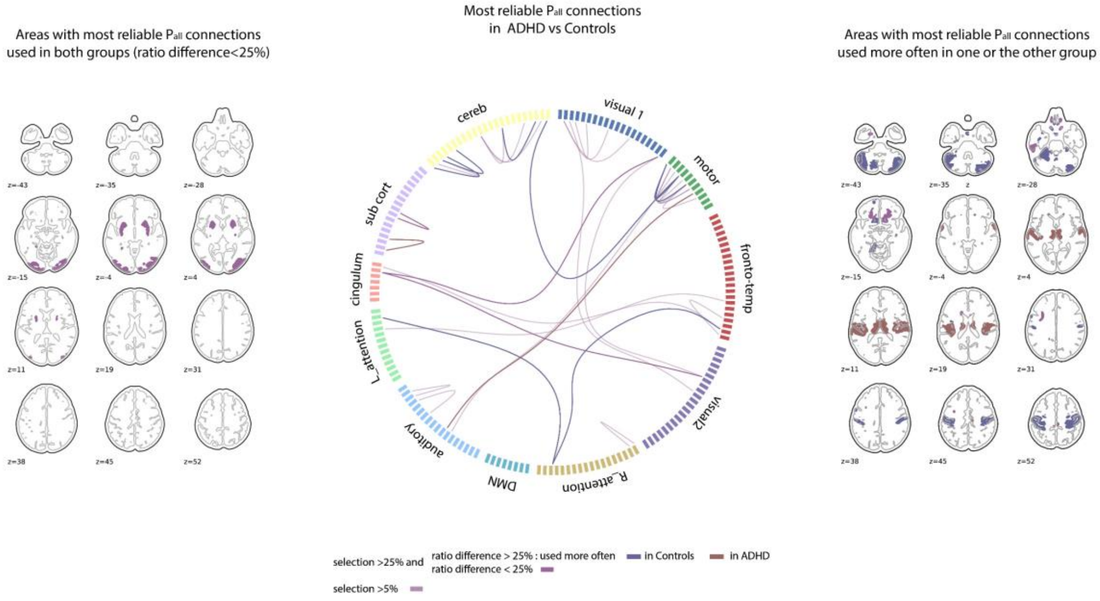
Areas with most reliable Pall connections used in both groups (ratio difference<25%)
Most reliable Pall connections in ADHD vs Controls
Areas with most reliable Pall connections used more often in one or the other group
cereb
visual 1
motor
fronto-temp
visual2
R_attention
DMN
auditory
L_attention
cingulum
sub cort
selection >25% and
ratio difference > 25% : used more often
in Controls
in ADHD
ratio difference < 25%
selection >5%

***Supplemental figure 6:*** *Representation of areas with most reproducible $P_{all}$ connections comparing groups. The brain slices on the left show areas with at least one connection used reproducibly (>50% of selection and ratio difference <25%) in both compared groups. The circular connectivity plot represents all connection selected >50% as $P_{all}$ in one of the two compare groups across bootstraps. Within these strongly reproducible $P_{all}$ connections, the one used most often in one group are represented with full color of the group (ratio difference >25%) (red for ADHD, blue for Controls, green for Siblígns. The brain slices on the right represent the associated areas to the connection used more often in one group. If an area has connections used in each group, its color is a blend of the compared group colors (purple for ADHD vs Control, yellow for ADHD vs Siblings and turquoise for Controls vs Siblings)*

Supplemental figure 6 shows the brain areas associated with the most reliably modulated $P_{all}$ connections in controls (above 50% across bootstraps). Within those connections, we further differentiated between those connections that were almost never modulated by ADHD (ratio of modulation percentage is above 50%) and those that were relatively more modulated by ADHD participants. Overall, controls most reliably modulated connections involving a widespread pattern of areas typically associated with reward and executive networks[19]. Observed areas included cerebellar, subcortical, motor and fronto-parietal areas.

#### Task potency differences

*Supplemental Table 4: test corresponding the differences between percentage of modulated edges between each group and their relative null distribution (see figure 1, 3 and supplemental figure 4).*

| | ADHD | | | Sibling | | | Control | | |
|---|---|---|---|---|---|---|---|---|---|
| | P-value | Mean(std) | Null distribution mean (std) | P-value | Mean(std) | Null distribution mean (std) | P-value | Mean(std) | Null distribution mean (std) |
| Sensitive edges | **0.02** | 0.402 (0.047) | 0.506 (0.041) | 0.107 | 0.445 (0.045) | 0.510 (0.040) | 0.3312 | 0.473 (0.044) | 0.501 (0.040) |
| $P_{all}$ edges | **0.04** | 0.139 (0.048) | 0.205 (0.036) | 0.107 | 0.163 (0.043) | 0.209 (0.036) | 0.3312 | 0.178 (0.045) | 0.195 (0.039) |
| $P_{mix}$ edges | 0.42 | 0.284 (0.055) | 0.293 (0.041) | 0.2457 | 0.255 (0.047) | 0.282 (0.039) | 0.3312 | 0.298 (0.051) | 0.320 (0.045) |
| $P_{single}$ edges | **0.04** | 0.577 (0.057) | 0.501 (0.041) | 0.129 | 0.581 (0.050) | 0.509 (0.041) | 0.3312 | 0.525 (0.052) | 0.485 (0.044) |

*Supplemental Table 5: Mean potency and standard deviation for each condition*

| | | Generic | | Unspecific | | Specific | |
|---|---|---|---|---|---|---|---|
| | | Shared | Unique | Shared | Unique | Shared | Unique |
| ADHD | p-value | 0.08 | 0.12 | 0.12 | **0.05** | 0.44 | **0.02** |
| | Mean (sd) | 2.056 (0.279) | 1.629 (0.180) | 1.648 (0.297) | 1.563 (0.142) | 1.232 (0.306) | 1.411 (0.0944) |
| | Null distribution mean (sd) | 1.792 (0.115) | 1.418 (0.178) | 1.420 (0.108) | 1.284 (0.093) | 1.193 (0.145) | 1.135 (0.048) |
| Siblings | p-value | 0.08 | 0.49 | 0.18 | 0.49 | 0.49 | 0.49 |
| | Mean (sd) | 2.160 (0.296) | 1.193 (0.264) | 1.684 (0.287) | 0.983 (0.179) | 1.307 (0.296) | 0.582 (0.113) |
| | Null distribution mean (sd) | 1.778 (0.113) | 1.165 (0.194) | 1.416 (0.107) | 0.945 (0.121) | 1.203 (0.132) | 0.576 (0.067) |
| Controls | p-value | 0.11 | 0.33 | 0.26 | 0.33 | 0.36 | 0.36 |
| | Mean (sd) | 2.133 (0.218) | 1.327 (0.277) | 1.708 (0.296) | 1.069 (0.198) | 1.183 (0.309) | 0.622 (0.119) |
| | Null distribution mean (sd) | 1.830 (0.122) | 1.177 (0.199) | 1.450 (0.107) | 0.942 (0.120) | 1.184 (0.176) | 0.580 (0.069) |

#### Task performance differences between groups

*Supplemental Table 6: independent t-test between task performances of the ADHD, Siblings and Controls. The strongest differences are observed in the stop signal reaction time variability for ADHD participants. No differences in performance is observed between Siblings of ADHD and the control participants.*

| Task performance | ADHD mean (sd) | Siblings mean (sd) | Control mean (sd) | ADHD vs Control p-val (t) | ADHD vs Siblings p-val (t) | Siblings vs Control p-val (t) |
|---|---|---|---|---|---|---|
| STOP - Stop signal reaction time | 265.5 (59.1) | 252.0 (48.7) | 255.5 (51.4) | 0.132 (-1.51) | 0.06 (1.92) | 0.609 (0.51) |
| STOP - reaction time variability | 111.1 (39.7) | 93.3 (37.0) | 84.0 (31.5) | **$1.3 \times 10^{-9}$ (-6.3)** | **0.0003 (3.6)** | **0.042 (-2.04)** |
| REWARD – reward and non-reward reaction time differences | 44.1 (50.1) | 34.2 (31.2) | 30.2 (32.1) | **0.008 (-2.7)** | 0.06 (1.8) | 0.35 (-0.9) |
| WM – working memory performance | 0.24 (0.04) | 0.24 (0.04) | 0.24 (0.04) | 0.975 (-0.05) | 0.968 (-0.08) | 0.951 (-0.12) |

#### Confounder effects

*Supplemental Figure 7: replication of the results presented in figures 1, 2, and SF4 in subsamples to evaluate possible confounder effects of medication (top graphs corresponds to the comparison of ADHD off (left) and on (right) medication with Siblings and Controls), gender (second row of graphs corresponds to the subsamples of only female (left) or only male (right) for each groups), scanner (third row of graph corresponds to the subsamples of participants acquired with the Siemens AVANTO (left) or the Siemens SONATA (right)), and co-morbidity (bottom graphs corresponds to the comparison of ADHD with another comorbid disorder (ODD, CD, Tourette or/and TIC) (left) or without comorbid disorders (right) compared to Siblings and Controls). Conclusions from the main analysis on $P_{all}$ edges hold for each condition, conclusions on $P_{single}$ edges shows less robustness as also concluded from figure 2. Differences are more strongly expressed in females than in males, however, the reduction of power does not enable to draw firm conclusions from these differences.*

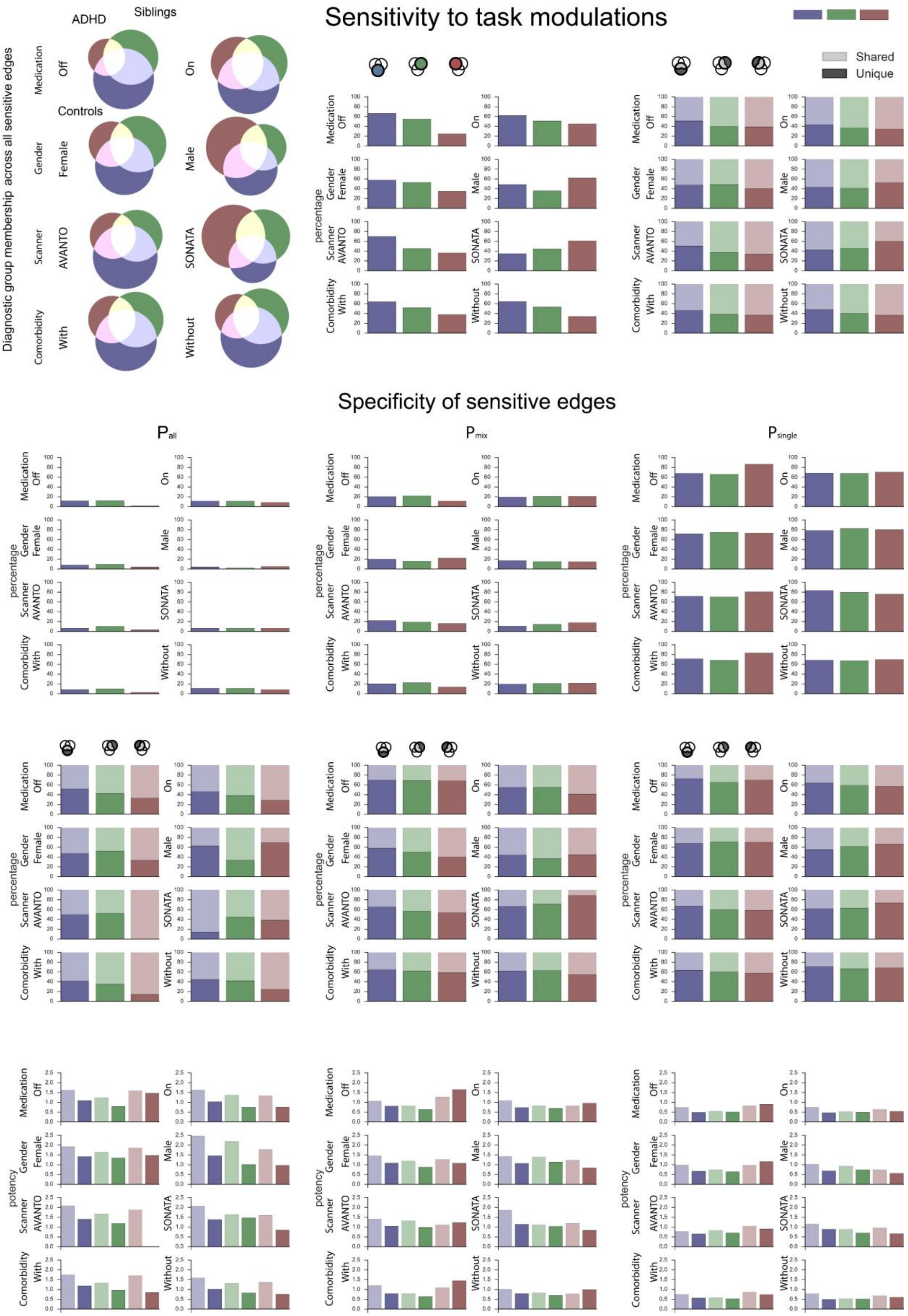
Sensitivity to task modulations
Specificity of sensitive edges
Diagnostic group membership across all sensitive edges
ADHD
Siblings
Controls
Shared
Unique
Medication
Off
On
Gender
Female
Male
Scanner
AVANTO
SONATA
Comorbidity
With
Without
percentage
potency
$P_{all}$
$P_{mix}$
$P_{single}$

*Supplemental Figure 8: replication of the results presented in SF4 in subsamples to evaluate possible confounder effects. We matched three groups on IQ, site of acquisition, age and gender using pymatch. We observe that the observed pattern from the main analysis using the bootstrapping is also observed using a matched sample. The variance estimation using a bootstrapping method is not here replicated as the number of subject in the matched sample is lower (min 30 max 42 participants per groups)*

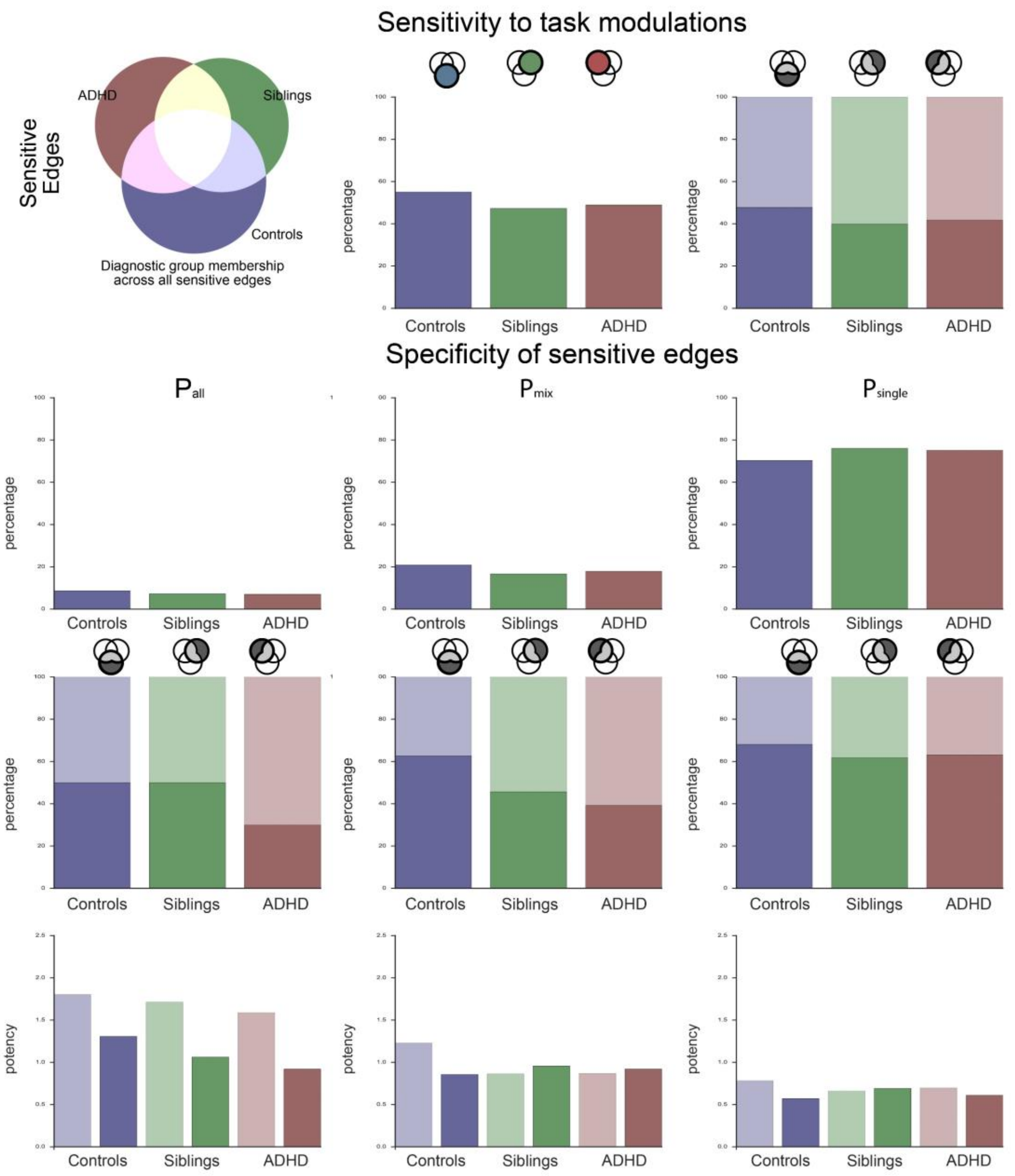
